# Optimization of a Cardiomyocyte Model Illuminates Role of Increased I_NaL_ in Repolarization Reserve

**DOI:** 10.1101/2023.08.31.555767

**Authors:** Kristin E Fullerton, Alexander P. Clark, Trine Krogh-Madsen, David J Christini

**Author notes:** Corresponding author: David Christini, SUNY Downstate Health Sciences University, 450 Clarkson Ave, Brooklyn, NY 11203.

## Abstract

Cardiac ion currents may compensate for each other when one is compromised by a congenital or drug-induced defect. Such redundancy contributes to a robust repolarization reserve that can prevent the development of lethal arrhythmias. Most efforts made to describe this phenomenon have quantified contributions by individual ion currents. However, it is important to understand the interplay between all major ion channel conductances, as repolarization reserve is dependent on the balance between all ion currents in a cardiomyocyte. Here, a genetic algorithm was designed to derive a profile of nine ion-channel conductances that optimizes repolarization reserve in a mathematical cardiomyocyte model. Repolarization reserve was quantified using a previously defined metric, repolarization reserve current, i.e., the minimum constant current to prevent normal action potential repolarization in a cell. The optimization improved repolarization reserve current by 77 % compared to baseline in a human adult ventricular myocyte model and increased resistance to arrhythmogenic insult. The optimized conductance profile was characterized by increased repolarizing current conductances, but also uncovered a previously unreported behavior by the late sodium current. Simulations demonstrated that upregulated late sodium increased action potential duration, without compromising repolarization reserve current. The finding was generalized to multiple models. Ultimately, this computational approach in which multiple currents were studied simultaneously illuminated mechanistic insights into how the metric’s magnitude could be increased, and allowed for the unexpected role of late sodium to be elucidated.

**NEW & NOTEWORTHY:** Genetic algorithms are typically used to fit models or extract desired parameters from data. Here, we utilize the tool to produce a ventricular cardiomyocyte model with increased repolarization reserve. Since arrhythmia mitigation is dependent on multiple cardiac ion-channel conductances, study using a comprehensive, unbiased, and systems-level approach is important. This use of this optimization strategy allowed us to find a robust profile that illuminated unexpected mechanistic determinants of key ion-channel conductances in repolarization reserve.

## 1 Introduction

Sudden cardiac death accounts for 20% of all deaths worldwide (Hayashi et al., 2015). Unfortunately, the development of a potentially fatal ventricular arrhythmia is complex and not well understood. Within each cardiomyocyte is a myriad of different ion channels and transporters expressed at varying densities which dictate a given cardiomyocyte’s action potential morphology (Marder and Goaillard, 2006; Weiss et al., 2012). Extremes in levels of some conductances due to congenital disease or drugs can lead to repolarization failure or early afterdepolarization (EAD) development, and susceptibility to arrhythmia (Fenichel et al., 2004; Kannankeril et al., 2010; Hancox et al., 2008). Even two seemingly healthy cardiomyocytes can differ in their robustness against arrhythmia when under the influence of a perturbation (Weiss et al., 2012; Ballouz et al., 2021; Miller et al., 2023). Unfortunately, a mechanistic understanding of repolarization abnormality development, and improvement of pharmacological mitigation strategies, is complicated by the many different ion-channel conductance profiles that can define a cardiomyocyte (Behr and Ensam, 2016). Therefore, defining the conductance values of key ion-channels, or quantifying a unique ion-channel conductance profile for each cell, can allow for a better understanding of the different degrees of robustness between cells.

One key manifestation of a cardiomyocyte’s conductance profile is its ability to repolarize in the presence of arrhythmogenic repolarization challenges (Roden, 2006). This ability can be quantified via “repolarization reserve current” (RRC), which is a measure of the robustness of a cell to effectively repolarize (Gaur et al., 2020) and can be used to assess inherent arrhythmia risk. It was developed recently to quantify “repolarization reserve” - a concept that describes the robustness of repolarization, meaning that a block in one ionic current might not cause the development of an EAD or repolarization failure because it can be compensated by other repolarzing currents (Roden, 1998; Varró and Baczkó, 2011). Therefore, repolarization reserve can be used to better understand cardiac arrhythmogenesis, and possibly inform drug development for mitigation.

While RRC is a valuable metric in arrhythmia risk assessment on the single-cell level, it lacks mechanistic transparency because it bundles all ion currents together (Gaur et al., 2020). Quantifying the corresponding ion-specific conductance profile would more fully describe a cardiomyocyte’s repolarization reserve and elucidate how variability in ion-channel densities translates to RRC values. While repolarization reserve has been studied, most efforts made to describe this phenomenon have quantified effects of individual ion channel conductances (Xiao et al., 2008; Jost et al., 2005; Varshneya et al., 2018; Silva and Rudy, 2005; Printemps et al., 2019; Ishihara et al., 2009; Jost et al., 2013; Biliczki et al., 2002). For example, the slow delayed rectifier potassium current (I_Ks_) (Xiao et al., 2008; Jost et al., 2005; Varshneya et al., 2018; Silva and Rudy, 2005) and the inward rectifier current (I_K1_) (Ishihara et al., 2009; Jost et al., 2013; Biliczki et al., 2002) have been independently demonstrated to provide the necessary repolarizing current to prevent proarrhythmia in the presence of rapid delayed rectifier potassium current (I_Kr_) block. However, such studies do not consider the values or correlations between other ion-channel conductances in a given cell’s profile. Research using a more comprehensive approach has been completed using methods such as a population of models (Britton et al., 2017; Miller et al., 2023) and multivariable regression (Sarkar and Sobie, 2011). These studies make conclusions about ion-channel variability and proarrythmic susceptibility, but only in a reduced repolarization reserve environment. Therefore, there is insufficient understanding of the mechanism to which the magnitude of RRC can be increased on a systems level. Since repolarization reserve, and arrhythmia mitigation, is dependent on the state of all cardiac ion channels, research using such an approach may be instructive.

The research presented here aims to derive a profile of nine ion-channel conductances that can improve the repolarization reserve of an adult ventricular cardiomyocyte model, using a systems-level approach. We used the optimized model system to identify unexpected mechanistic determinants of the importance and behavior of key ion-channel conductances. Results show that the derived profile is more robust than the baseline model through its ability to attenuate proarrhythmic dynamics. This work is valuable because it provides a systems-level understanding of increased repolarization reserve, which has the potential to contribute to research in arrhythmia mitigation.

## 2 Methods

### 2.1 Computational Modeling

As our baseline model, we used the human-based endocardial Tomek et al. (Tomek et al., 2019) (ToR-ORd) model which represents the electrophysiology of a whole-cell adult ventricular cardiomyocyte. It also includes a detailed representation of intracellular calcium handling. Results were validated in the Grandi et al. (Grandi et al., 2010) (GB) endocardial model, which was updated to include the late sodium current, I_NaL_ (specific details in the Supplementary Material). Both models were paced at 1 Hz; the ToR-ORd model and GB models were stimulated at −53 A/F for 1 ms and −9.5 A/F for 5 ms, respectively. Myokit was used for all cardiomyocyte model simulations (Clerx et al., 2016).

### 2.2 Genetic Algorithm Design

A genetic algorithm is a method used to search for solutions through a process inspired by natural selection and has been used in the field to build models or extract desired parameters from data (Clark et al., 2022; Tomek et al., 2019; Bot et al., 2012; Groenendaal et al., 2015; Syed et al., 2005; Kherlopian et al., 2011). Through this technique, a population of individuals evolve towards an optimum in an iterative process. All individual models in an iteration, or generation, are assigned an error score based on a user-defined cost function. The individuals with the lowest error are then selected, mated to generate a new population, and mutated to ensure diversity for the next generation. This process continues for a set number of generations or until a minimum is reached.

We used a genetic algorithm, as implemented in Python and the DEAP package (Fortin et al., 2012), as our optimization method. The optimization allowed changes to the maximal conductance or flux of nine currents: I_K1_, I_Kr_, I_Ks_, I_NaL_, sodium (I_Na_), L-type calcium (I_CaL_), transient outward (I_to_), sodium-calcium exchanger (I_NCX_), and sodium-potassium pump (I_NaK_). Each individual in the genetic algorithm is a unique profile of nine key current conductance or maximal flux scaling factors (we will refer to these collectively as conductances). Scaling factors were sampled from a log-normal distribution within an allowed range of 33% to 300% of baseline. We used a population size of 200 individuals and ran the algorithm over 100 generations. Since genetic algorithms are innately stochastic, the optimization was run for 8 trials.

The cost function to be minimized during the optimization was developed to ensure that the algorithm optimizes the model to both (1) have an increased RRC magnitude and (2) be physiologic (Figure 1.). Specifically, it consists of two terms that penalize against deviations of the action potential (AP) and several calcium transient markers from physiological ranges. A third term penalizes against low RRC values. This cost function (Equation 4 below) is evaluated for each individual (*i*) during the evolution:

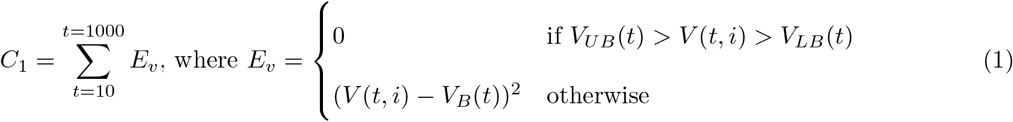

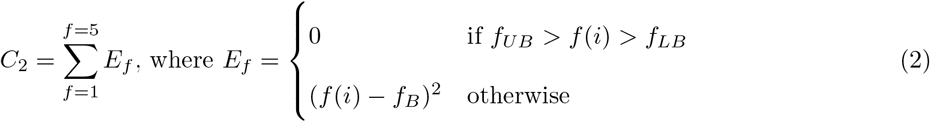

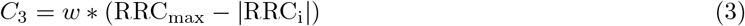

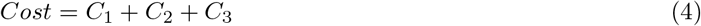

**Figure 1:**
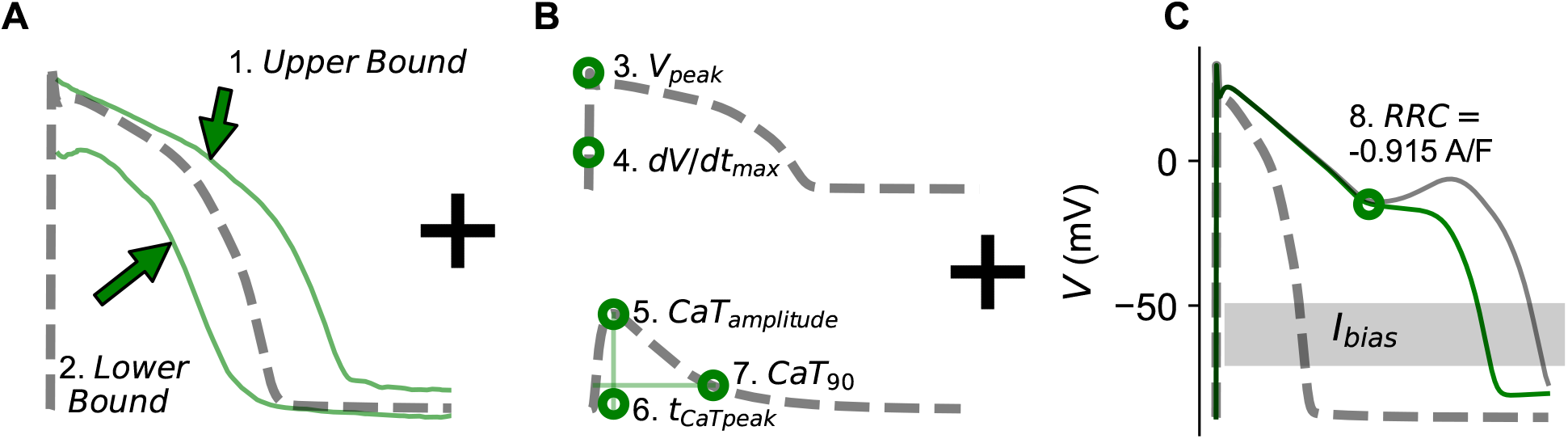
Graphical representation of the cost function designed for genetic algorithm optimization. A. Upper and lower action potential bounds to assess action potential morphology point by point after upstroke. The morphology error is zero when the voltage at every time point is within the upper and lower bounds (Tomek et al., 2019; Britton et al., 2017). B. Features calculated to assess action potential upstroke and calcium transient morphology. To calculate the feature error for a given individual, the peak voltage (*V*_*peak*_), maximal rate of rise (*dV/dt*_*max*_), calcium transient amplitude (*CaT*_*amp*_), calcium transient duration at 90% (*CaT*_90_), and the time to reach peak amplitude (*tCaT*_*peak*_) are calculated. If a given feature is within its upper and lower bounds, the error for that feature term is zero. C. Repolarization reserve current (RRC) is calculated by injecting a constant depolarizing bias current, I_bias_, in small increments until a repolarization abnormality is apparent. I_bias_ current was injected 4 ms after upstroke until the end of the action potential cycle length (grey bar denotes the time duration of the I_bias_ injection). The RRC value is the minimum I_bias_ current value at which the action potential can repolarize normally. I_bias_ = 0 A/F for the dashed grey line; I_bias_ = −0.915 A/F for the solid green line; and I_bias_ = −0.938 for the solid grey line.

Equation 1 assigns a penalty if the simulated action potential (after its upstroke) is not physiologic (Figure 1A). Similarly, Equation 2 assigns a penalty based on the values of five action potential upstroke and calcium transient features (*f*) (Figure 1B and Table 1). Therefore, *C*_1_ and *C*_2_ together penalize against individuals that are not physiologically relevant. In both, an error of 0 is given if the voltage or feature value is within bounds; otherwise, the error is the squared difference between the given value and that of the baseline ToR-ORd model. *V*_*B*_(*t*) and *f*_*B*_ represent a given voltage and feature value for the baseline ToR-ORd model, respectively. The errors are then summed for each time point in the cycle length of an action potential (*C*_1_), or for every feature (*C*_2_).

**Table 1:** Upper bound, lower bound, and baseline feature values that were used to calculate the cost for every individual in the genetic algorithm optimization. The features include: *V*_*peak*_, *dV/dt*_*max*_, *CaT*_*amp*_, *tCaT*_*peak*_, and *CaT*_90_. Upper and lower bounds were chosen by selecting the maximum and minimum value, respectively, of each experimentally recorded biomarker reported in the literature (Passini et al., 2017; Britton et al., 2013; O’Hara et al., 2011; Coppini et al., 2013; Tomek et al., 2019; Beuckelmann et al., 1992; Krogh-Madsen et al., 2017). The middle column titled ToR-ORd model represents the feature value of the baseline ToR-ORd model.

| Feature ($f$) | Lower Bound ($LB$) | ToR-ORd Model ($B$) | Upper Bound ($UB$) |
| --- | --- | --- | --- |
| $V_{peak}(mV)$ | 10 | 33 | 55 |
| $dV/dt_{max}(mV/ms)$ | 100 | 347 | 1000 |
| $CaT_{amp}(\mu M)$ | 0.3 | 0.312 | 0.8 |
| $tCaT_{peak}(ms)$ | 40 | 58 | 60 |
| $CaT_{90}(ms)$ | 350 | 467 | 500 |

Equation 3 penalizes against individuals that have a low RRC. The RRC for a given individual (RRC_i_) is calculated and subtracted from RRC_max_ = 3 A/F which is an arbitrary and extreme upper bound to evaluate RRC magnitude. That difference is then multiplied by a weight of *w* = 20000 to account for the inherent differences in variable values.

### 2.3 Biomarkers and Physiologic Constraints

The features used to assess physiologic relevancy for the action potential upstroke and calcium transient are listed in Table 1 and include: peak membrane potential (*V*_*peak*_), action potential maximal rate of rise (*dV/dt*_*max*_), calcium transient amplitude (*CaT*_*amp*_), time at which the calcium transient is maximal (*tCaT*_*peak*_), and calcium transient duration at 90% decay (*CaT*_90_). The action potential upstroke feature (*V*_*peak*_ and *dV/dt*_*max*_) bounds were based on a population-of-models study (Passini et al., 2017) and human action potential recordings (Britton et al., 2013; O’Hara et al., 2011). The calcium transient feature bounds were informed by various human data sources (Coppini et al., 2013; O’Hara et al., 2011; Tomek et al., 2019; Beuckelmann et al., 1992; Krogh-Madsen et al., 2017). The lower and upper bounds used in Equation 1 (*V*_*LB*_(*t*) and *V*_*UB*_(*t*), respectively) were the 10th and 90th percentiles of the Szeged-ORd experimental action potential dataset (Tomek et al., 2019; Britton et al., 2017).

Equations 1 and 2 were calculated for five consecutive action potentials (after pre-pacing each individual for 600 beats) and summed to arrive at final values for *C*_1_ and *C*_2_. Preliminary simulations with a 10 model data set had demonstrated that action potential duration at 90% repolarization (APD_90_) is very consistent after 100 beats, with beat-to-beat differences less than 1 ms for action potentials with normal repolarization. Therefore, the 605 beat simulation number was chosen to ensure steady-state.

### 2.4 RRC Calculation

As previously described (Gaur et al., 2020), RRC is determined by finding the minimum constant current, I_bias_, required to prevent normal cardiomyocyte repolarization (Figure 1C). The typical protocol includes injection of a constant inward (i.e., negative) current that increases in magnitude, spaced by a few baseline beats (I_bias_=0), until a repolarization abnormality develops. RRC is then taken as the most negative current value injected that allowed for normal repolarization. More negative current values indicate increased repolarization reserve.

As in Gaur et al. (2020), the depolarizing I_bias_ current was injected 4 ms after upstroke until the end of the action potential cycle length. Action potentials with current injections were spaced by 4 beats without current injection. Repolarization abnormality was defined as an action potential presenting an EAD, repolarization failure, or instability of phase four of the action potential. An EAD was taken to occur if the slope of two consecutive points after action potential peak (*t >* 100 ms) was positive. A repolarization failure was defined if an action potential did not repolarize below −70 mV. In addition, if phase four had a positive slope greater than 0.01 mV/ms, the action potential was deemed as unstable.

For computational efficiency, we utilized a binary search technique for automatic RRC calculation, and continued the search until the difference between the sizes of the currents leading to normal and abnormal repolarization became less than 0.025 A/F. This RRC protocol was done for each individual immediately following the 605 baseline beats.

### 2.5 Sensitivity Analyses

A local sensitivity analysis was performed for the baseline and optimized ToR-ORd models to evaluate the effect of ion-channel conductance perturbations on the cost function (Groenendaal et al., 2015). Each ionchannel conductance was varied by 30% and 300% of their baseline values and the sensitivity computed as the change in each component of the cost function (i.e., Equations 1, 2, and 3) from baseline. This approach was used to assess *C*_3_ (RRC) sensitivity to each ion-channel conductance in the GB model as well.

### 2.6 Statistical Methods

To analyze pair-wise relationships between ion-channel conductances Spearman correlation was used. One, two, or three stars in Figure 6 denote a p-value of 0.05, 0.01, and 0.001, respectively.

## 3 Results

### 3.1 Genetic algorithm design for optimization of a conductance profile with improved RRC

We used a genetic algorithm and *in silico* cardiomyocyte model to find an RRC-improving ion-channel conductance profile. RRC for each profile was determined as the maximal constant inward current value that can be injected into a cell that allows for normal repolarization (Gaur et al., 2020) as described in the Methods. For example, the RRC for the baseline ToR-ORd model is −0.915 A/F (green, solid) as the next current injection of −0.938 A/F (grey, solid) caused an EAD (Figure 1C). We ran 8 optimizations to account for the inherent stochastic nature of genetic algorithms. All simulations converged within 100 generations (Figure 2A).

**Figure 2:**
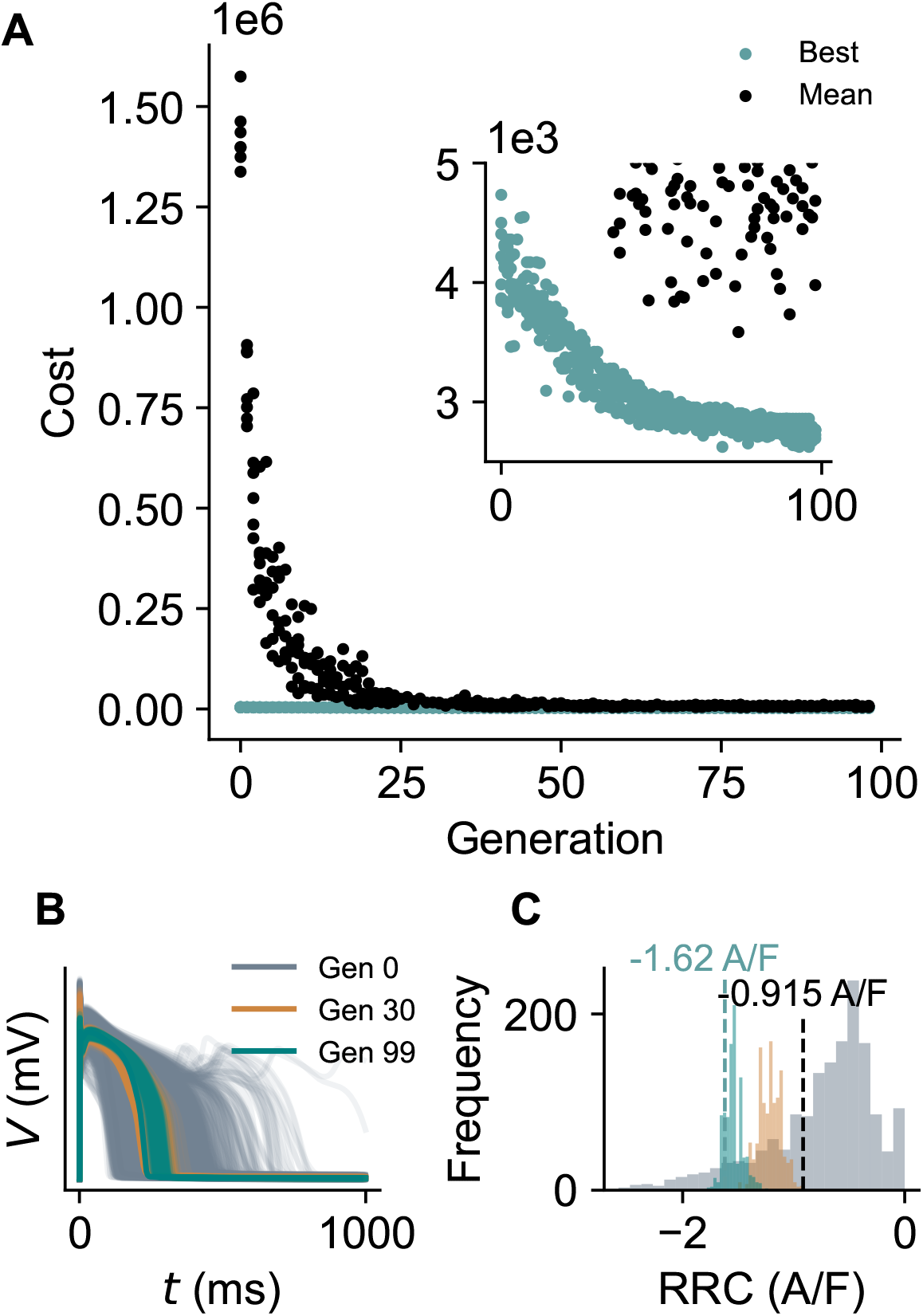
A genetic algorithm that optimizes for RRC while ensuring physiologic morphology. A. Optimization progression of 200 individuals over 100 generations. A low cost score represents a physiologic individual with a high RRC. The average cost for each generation for 8 trials is plotted in black. The cost of the best individual for each generation is plotted in teal. B. Action potentials from each individual in generations 0 (grey), 30 (orange), and 99 (teal). C. RRC distributions for all individuals of all trials for generations 0 (grey), 30 (orange), and 99 (teal). The black and teal dashed lines indicate the baseline RRC and mean RRC for the best 220 optimized models, respectively. The optimized model improved RRC by 77% compared to the baseline ToR-ORd model.

During the optimization, action potentials (AP) converge to within the physiological range (Figure 2B) and RRC improves (Figure 2C). Indeed, while even the best individuals in the final generation had a non-zero cost (Figure 2A), they all produce physiologic APs and calcium transients as well as improved RRC. Therefore, the non-zero cost value after convergence is entirely due to the RRC term (*C*_3_) in Equation 4 (Figure S1) which is expected as the reference RRC_max_ value of 0.3 pA/pF is an extreme upper bound. Note also that while the initial generation had individuals with even better RRC values than those found after 100 generations, these individuals generated unphysiologic APs and/or calcium transients.

### 3.2 The optimized models are more robust against proarrythmia than the baseline ToR-ORd model

The genetic algorithm was successful in finding multiple solutions that attenuate abnormal repolarization by increasing the model’s RRC. The best 220 individuals (i.e., those with *Cost <* 2800) were selected to visualize the ion-channel conductance profile derived (Figure 3A). Conductances of outward currents were increased and those of inward currents decreased, with the notable exception of I_NaL_. The resulting APs and calcium transients prove to be within physiological bounds as demonstrated in Figures 3B and 3C and have an average RRC of −1.62 A/F - a 77% improvement compared to the baseline ToR-ORd model.

**Figure 3:**
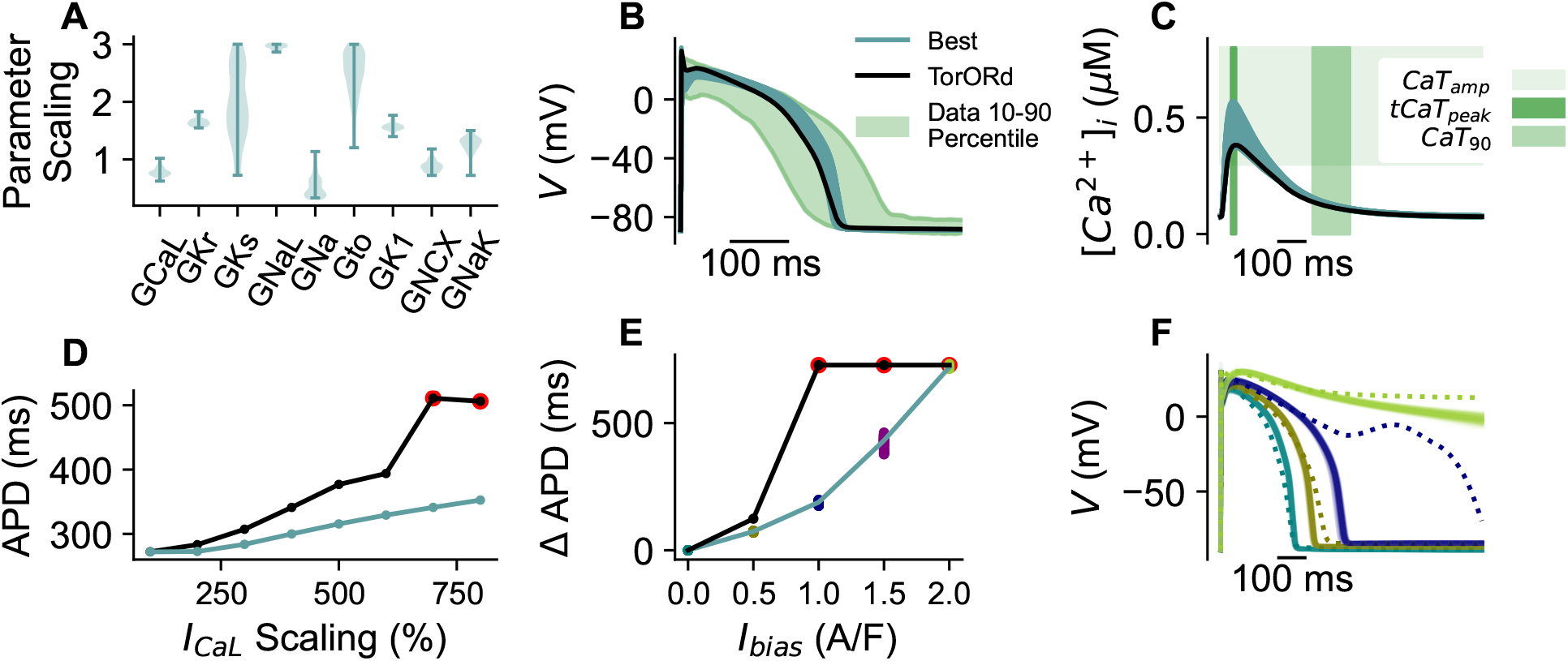
The best individuals found by the genetic algorithm are physiologic and allow for increased resistance to proarrythmic substrates. A. Scaling of conductance values for the best individuals (those with a cost lower than 2800; n = 220). The scaling factor is relative to the baseline ToR-ORd model (parameter scaling is 1). B. Action potential data for the best 220 individuals. The traces are compared to 10-90 percentile of experimental data (shaded regions) as detailed in the Methods (Britton et al., 2017; Tomek et al., 2019). C. Calcium transient morphology for each of the 220 best individuals compared to the baseline Tor-ORd model (black). D. APD_90_ values after perturbing the baseline ToR-ORd (black) and a representative optimized model (teal) by various *I*_*CaL*_ perturbations. The red circles indicate simulations in which a repolarization abnormality (EAD or repolarization failure) occurred. E. Change in APD_90_ between the baseline morphology of every best individual and the following I_bias_: 0, 0.5, 1.0, 1.5, 2.0 A/F. The same analysis was completed for the baseline ToR-ORd model (black). F. Action potential traces of simulations used to generate Figure 3E. The dotted and solid lines represent output from the baseline and optimized models, respectively, at I_bias_ = 0, 0.5, 1.0, and 2.0 A/F.

The optimized models also had increased resistance to proarrhythmia. APD_90_ was evaluated after various I_CaL_ perturbations in Figure 3D for both the baseline ToR-ORd (black) and a representative model from the 220 best GA individuals (teal; chosen randomly). It is clear that the APD_90_ for the optimized model is smaller at every I_CaL_ perturbation compared to that of the baseline model. This also demonstrates the ability of the optimized model to protect against proarrythmia. Unlike the baseline model, the optimized model was able to repolarize after 700% and 800% I_CaL_ scaling. This concept was also tested with I_Kr_ block (Figure S3A). While APD_90_ increases the more I_Kr_ is blocked, no repolarization abnormalities were evident, even at 99% block of I_Kr_ neither for the baseline nor the optimized model. This makes it challenging to evaluate the robustness of the optimized model to I_Kr_ perturbations. To further evaluate, and initiate repolarization abnormalities, we simulated block of both I_Kr_ and the background potassium channel (I_Kb_) (Figure S3B). With 40% I_Kb_ and 99% I_Kr_ block, repolarization failure was clear in the baseline model but not in the optimized model.

The impact of ion-channel block on repolarization is dependent on conductance scalings which can make an exact comparison between the baseline and the optimized models difficult. Therefore, lastly, the robustness of each optimized model was quantified by measuring the change in APD_90_ at various I_bias_ values (Figure 3E). The optimized models exhibited much less AP prolongation than the baseline model, indicating that all the profiles had increased resistance to potentially proarrhythmic repolarization delay. Figure 3F plots all optimized models (solid) and the baseline model (dotted) when perturbed by various I_bias_ stimuli: 0, −0.5, −1.0, and −2.0 A/F. When an I_bias_ value of −1.0 A/F is injected into all models, it is clear that the baseline model (navy, dotted) develops an EAD while the optimized models (navy, solid) do not. When I_bias_ is set to −2.0 A/F, both the baseline (green, dotted) and optimized models (green, solid) fail to repolarize. Therefore, Figures 3D, 3E, 3F, and S3 demonstrate the robustness of the optimized individuals compared to the baseline ToR-ORd model.

### 3.3 I_NaL_ prevents against excessive action potential shortening while maintaining improved RRC

As mentioned above, the ion-channel conductance profile found by the genetic algorithm increases outward currents and decreases inward currents, except I_NaL_ (Figure 3A). Indeed, the scaling of I_NaL_ hit its maximal allowed value of 3. This was unexpected as increased I_NaL_ is known to decrease repolarization reserve (Moreno and Clancy, 2012; Coppini et al., 2013; Ju et al., 1996; Trenor et al., 2012; Undrovinas et al., 2006; Bai et al., 2014; Hegyi et al., 2019; Valdivia et al., 2005). Therefore, steps were taken to elucidate the role of increased I_NaL_ in the model of increased arrhythmia resistance. I_CaL_, an inward current from the optimal ion-channel conductance profile with expected decreased conductance, was used as a reference.

Figure 4A demonstrates that APD_90_ is very sensitive to I_NaL_, but not as much to I_CaL_. Removal of I_NaL_ from the optimized ToR-ORd model caused a 51% reduction in APD_90_ (and AP morphology that is outside of the physiologic bounds), compared to a 9% reduction upon removal of I_CaL_.

**Figure 4:**
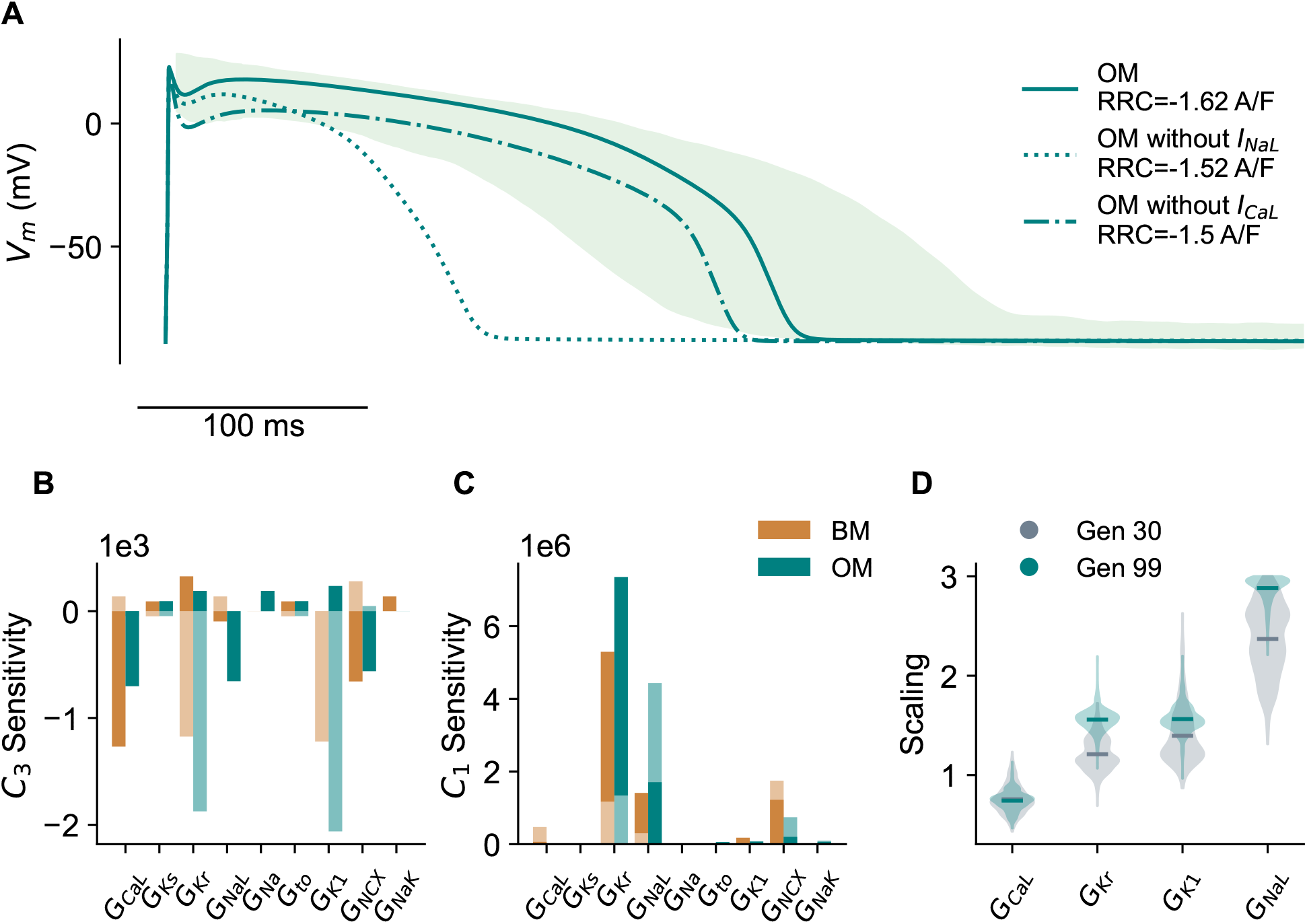
I_NaL_ plays a key role to balance high RRC and physiologic APD_90_. A. AP morphology for a random model optimized by the GA (optimized model) under various conditions: 0% I_NaL_ and 0% I_CaL_. B. *C*_3_. C. *C*_1_ sensitivities to the major ion-channel conductances for both the baseline ToR-ORd (BM) and optimized models (OM). The darker and lighter bars indicate sensitivity of a given cost term (i.e. *C*_1_, *C*_2_) to an enhanced (darker) or blocked (lighter) conductance. D. Variance and mean of conductances key in GA optimization using a population of the best 100 individuals at generation 30 and 99.

A sensitivity analysis demonstrates that *C*_3_ (cost function for RRC optimization) is most sensitive to I_K1_ for the optimized model, and I_CaL_ for the baseline model (Figure 4B). For example, a decrease in I_K1_ will decrease RRC more than any other ion channel in the optimized model. In addition, *C*_1_ (AP morphology optimization) is very sensitive to I_NaL_ for the optimized model, but not as much for the baseline ToR-ORd model. This demonstrates that I_NaL_ is important for a physiologic action potential morphology to be maintained in a model with increased RRC magnitude.

Furthermore, distributions of four ion-channel conductances from the best 100 individuals of generations 30 and 99 are plotted in Figure 4D. I_CaL_ does not change in mean or variance from generation 30 to 99. Conversely, I_NaL_, and the two most important ion-channel conductances in optimizing RRC (I_Kr_ and I_K1_), continue to increase from generation 30 to generation 99. As demonstrated in Figure 2, physiologic constraints *C*_1_ and *C*_2_ are largely met by generation 30 (Figure 2B), but RRC continues to improve until generation 99 (Figure 2C). Therefore, Figure 4 demonstrates that I_NaL_ increases to allow the GA to further optimize RRC. Specifically, the increase in inward current is necessary to prevent extreme APD shortening that would result from increases in outward I_Kr_ and I_K1_ made by the GA to increase RRC magnitude.

### 3.4 I_CaL_ cannot increase APD without decreasing RRC due to its correlation with I_NCX_

While an increase in inward current is necessary to prevent the extreme APD shortening that would result from increases in outward current to increase RRC, it is intriguing why the genetic algorithm chose I_NaL_ over another inward current, like I_CaL_, for example.

To better understand I_CaL_ and I_NaL_, and their relationship with APD_90_ and RRC, these two ion-channel conductances were studied individually in the baseline ToR-ORd model. Figure 5 analyzes the effects of increases in I_CaL_ and I_NaL_ in the baseline model with improved RRC. However, instead of increasing RRC magnitude using ion-channel conductances as was done in the optimized model, the size of RRC was increased to −1.78 A/F by a constant outward bias current (I_bias_). As expected, the increase in outward bias current (BM+) shortened the action potential to an unphysiologically short value (Figure 5A). Attempting to rescue this phenotype by increasing I_CaL_ 3-fold (BM-I_CaL_), resulted in AP morphology with an unphysiologically depolarized dome and APD still falling outside the target range (too short), while RRC is substantially reduced in size (from −1.78 A/F to −1.1 A/F). Further, the increased I_CaL_ leads to an unphysiologically large *CaT*_*amp*_ (Figure 5B). In contrast, amplifying I_NaL_ (also 3-fold, BM-I_NaL_) led to normal AP dome, on-target APD, and physiologic *CaT*_*amp*_, while barely decreasing RRC magnitude (to −1.69 A/F).

**Figure 5:**
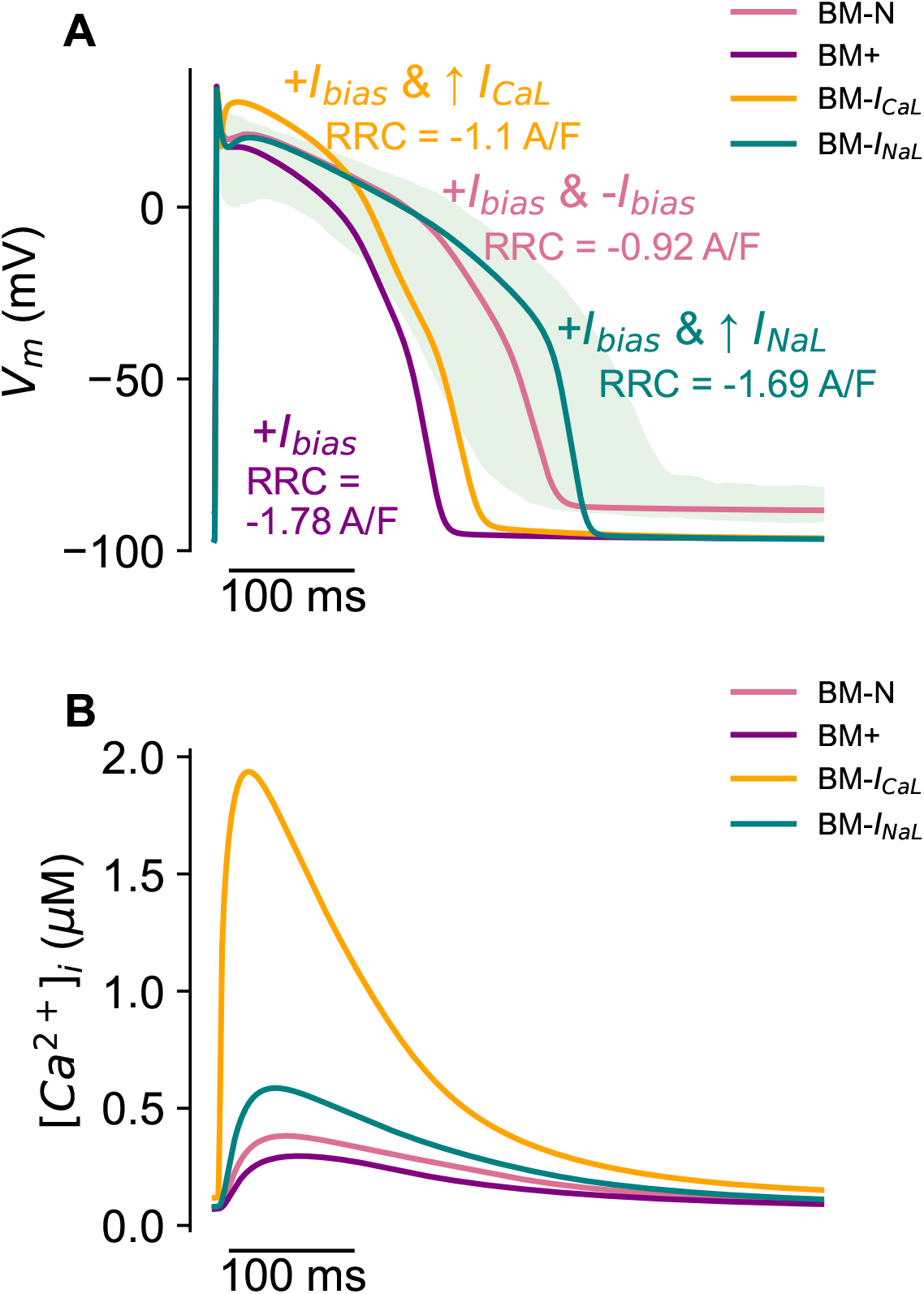
I_CaL_ cannot increase APD_90_ without decreasing RRC. A. ToR-ORd action potential traces under the following conditions: −0.8 A/F bias inward and 0.8 A/F bias outward current (BM-N), 0.8 A/F bias outward (BM+), 0.8 A/F bias outward and 300% I_CaL_ (BM-I_CaL_), 0.8 A/F bias outward and 300% I_NaL_ (BM-I_NaL_). B. Calcium transient traces for BM-N, BM+, BM-I_CaL_, and BM-I_NaL_.

To investigate how this finding translates to the optimized models, a correlation analysis was performed between the conductances of the 220 best GA individuals. Figure 6A demonstrates that I_CaL_ is correlated with I_NCX_ while I_NaL_ is not. Figure 6B plots I_NCX_ over time for BM+, BM-I_CaL_, and BM-I_NaL_ of Figure 5. It is clear that inward I_NCX_ is much increased when I_CaL_ is enhanced 3-fold (BM-I_CaL_, Figure 6B) compared to the case in which I_NaL_ is similarly augmented (BM-I_NaL_). The increase in inward I_NCX_ during phase 3 of the action potential possibly explains why RRC magnitude is increased upon I_CaL_ enhancement (BM-I_CaL_, Figure 5). When I_CaL_ conductance is increased, I_NCX_ must also increase to remove the excess calcium from the cell. In turn, the increase in inward current due to I_NCX_ decreases the repolarization reserve. In addition to I_CaL_ being an inward current, this result suggests that an increase in I_CaL_ decreases RRC due to its correlation with I_NCX_. Ultimately, these results allowed us to determine that (1) I_CaL_ cannot increase APD_90_ without decreasing RRC; and (2) I_NaL_ plays a key role in fine tuning high RRC and physiologic APD_90_.

**Figure 6:**
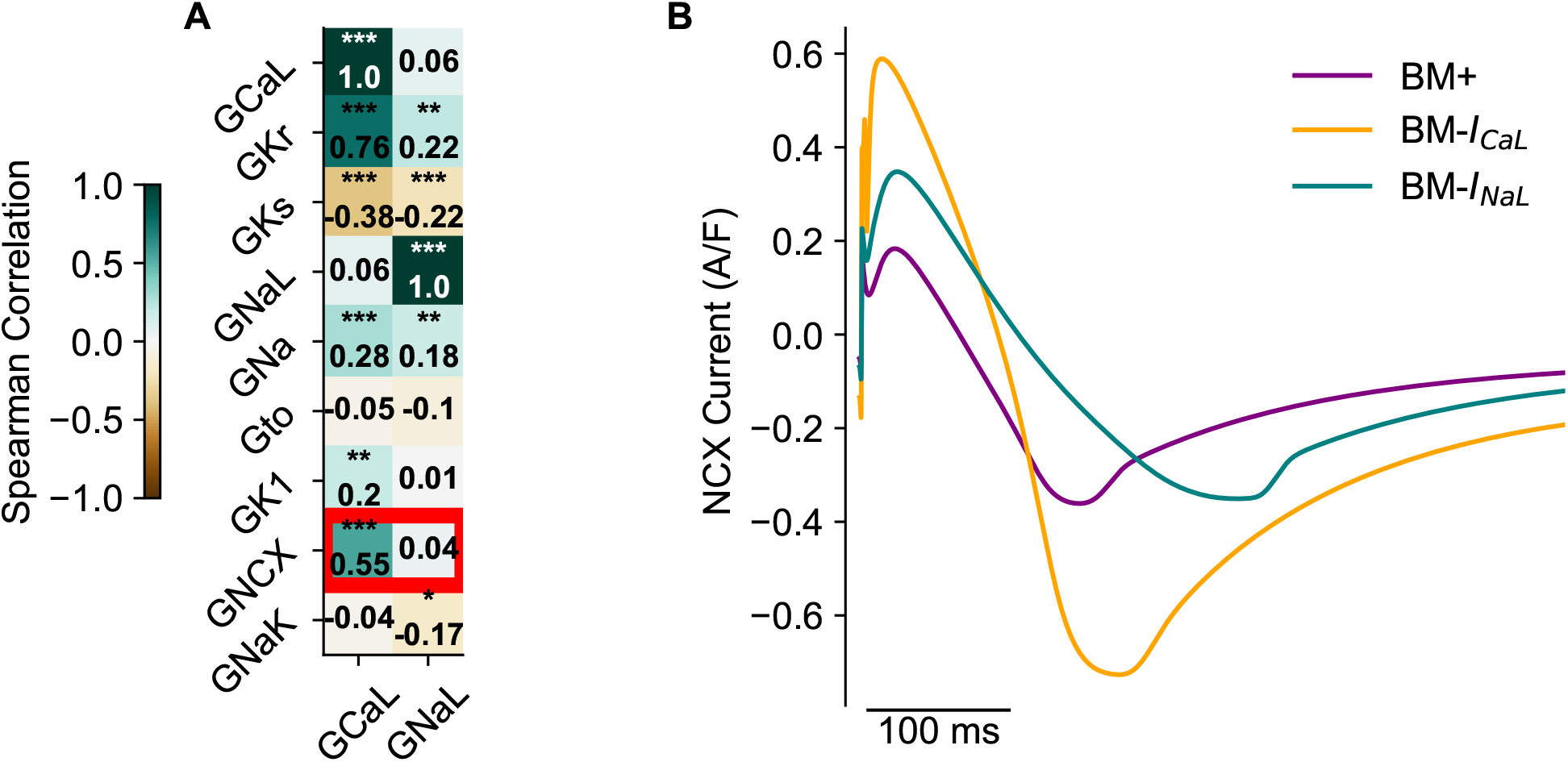
Unlike I_NaL_, I_CaL_ is correlated with I_NCX_ which prohibits it from increasing APD_90_ without decreasing RRC. A. Spearman correlations between G_CaL_ and G_NaL_ and all GA parameters for the population of 220 solutions with improved RRC. The red box highlights a strong correlation between G_NCX_ and G_CaL_ that is not present between G_NCX_ and G_NaL_. B. I_NCX_ current traces for BM+ (0.8 A/F bias outward), BM-I_CaL_ (0.8 A/F bias outward and 300% I_CaL_), and BM-I_NaL_ (0.8 A/F bias outward and 300% I_NaL_).

### 3.5 Addition of I_NaL_ to the GB model allows for RRC to be improved

As shown in Figure 5, I_NaL_ plays an important role in improving RRC while allowing APD to remain physiologic. To ensure that this dynamic is not specific to the ToR-ORd model, the relationship was studied in the GB model. The baseline GB model has a much reduced RRC compared to the ToR-ORd model and does not include I_NaL_ (Figure 7).

**Figure 7:**
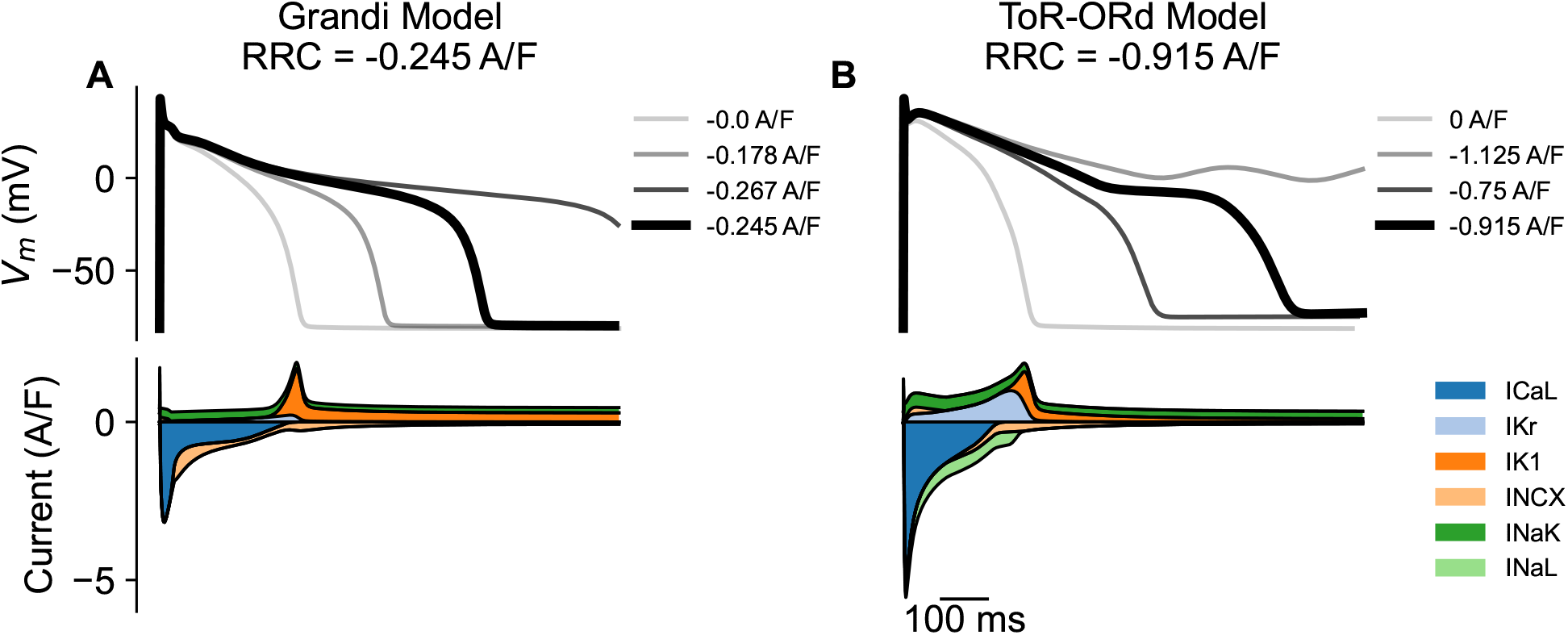
The RRC of the baseline GB model - which does not include I_NaL_ - is decreased 73% compared to ToR-ORd. A. RRC calculation (upper) and current densities (lower) for the baseline (A.) GB and (B.) ToR-ORd models.

In Figure 8A, a constant bias outward current was used to increase RRC magnitude of the baseline GB model to −1.14 A/F (GB+). When I_CaL_ was used to rescue the extreme shortening by GB+, physiologic APD was achieved, but RRC size dropped significantly (GBM-I_CaL_).

**Figure 8:**
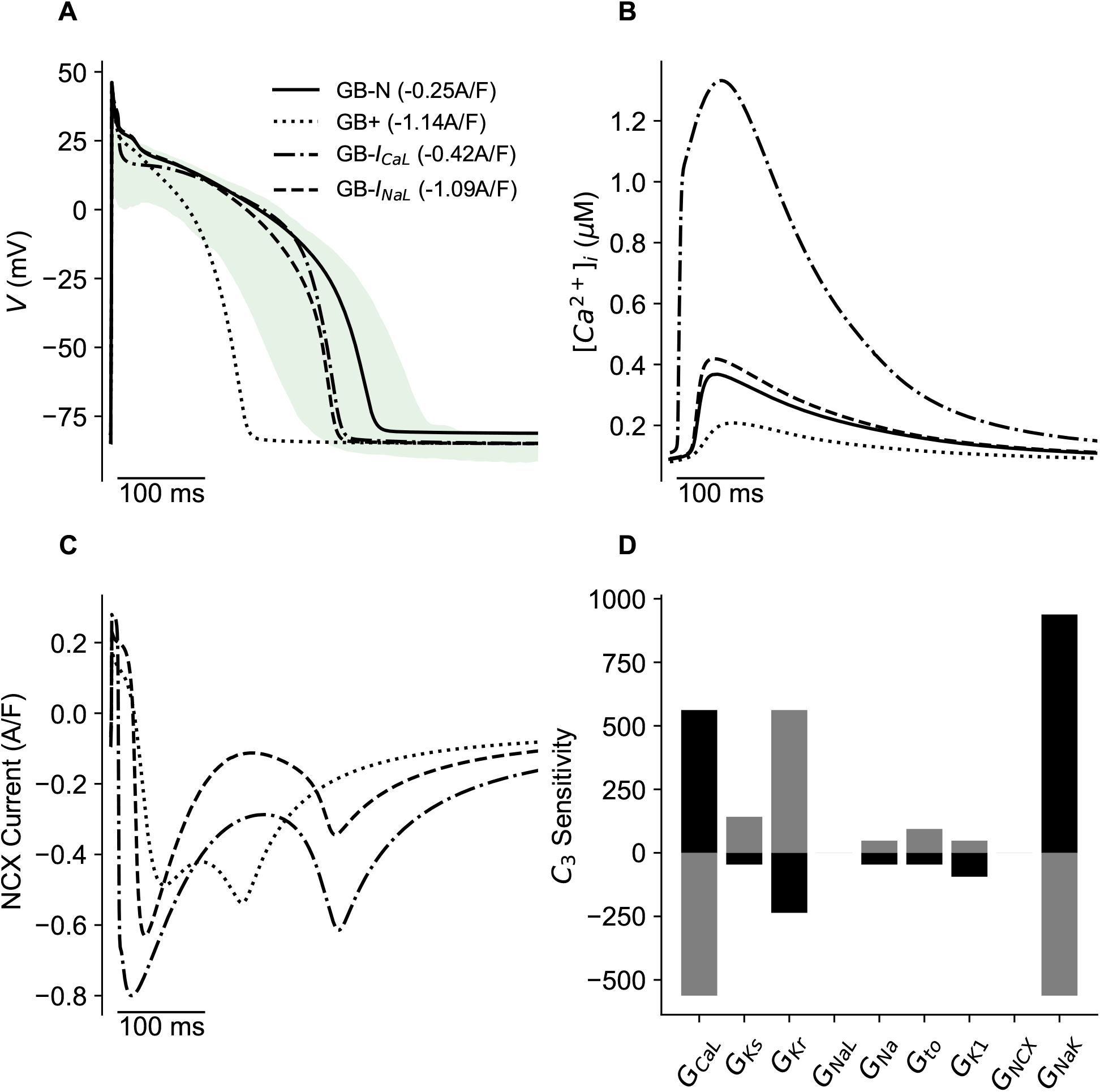
Addition of I_NaL_ to the GB model allows for RRC to be improved. A. GB action potential traces under the following conditions: −0.8 A/F inward bias and 0.8 A/F outward bias current (GB-N), 0.8 A/F outward bias (GB+), 0.8 A/F outward bias and 300% I_CaL_ (GB-I_CaL_), 0.8 A/F outward bias and 100% I_NaL_ (GB-I_NaL_). The RRC value for each action potential trace is in parenthesis. B. Calcium transient traces and C. I_NCX_ current for GB+, GB-I_CaL_, and GB-I_NaL_. D. RRC sensitivities to the major ion-channel conductances. The grey and black colored bars indicated augmentation and block of each conductance, respectively.

In addition, the upregulation of I_CaL_ caused *CaT*_*amp*_ to increase to a non-physiologic value (Figure 8B). To analyze the relationship between I_NaL_ and RRC in the GB Model, the ToR-ORd formulation of the current was added as described in the Supplementary Methods. With I_NaL_, APD was increased to a physiologic value and RRC magnitude decreased by only 0.05 A/F (GBM-I_NaL_). This confirmed that the relationship between I_NaL_ and RRC is not only specific to the ToR-ORd model. The mechanism to which RRC is decreased upon enhancement of I_CaL_ appears to be due to increased intracellular calcium. Figure 8C suggests that an increase in inward current by I_NCX_ decreased RRC size when APD is increased by I_CaL_ due to the large increase in intracellular calcium. Figure 8D further demonstrates this relationship as it shows I_CaL_ is the second most important ion-channel in RRC sensitivity. Unlike ToR-ORd in which RRC is most sensitive to I_CaL_ and I_K1_, RRC is most sensitive to I_NaK_ in the GB model. Despite the differences in RRC sensitivity between the ToR-ORd and GB models, the ability of I_NaL_ to rescue APD shortening by increased RRC is consistent.

## 4 Discussion

### 4.1 Main Findings

Here, we developed a genetic algorithm pipeline that searches for physiologically relevant ion-channel conductance profiles with improved RRC (Figure 2). This unbiased systems-level approach allowed for the derivation of a profile that improves RRC of the ToR-ORd model by 77%. It also illuminated unexpected insights regarding the mechanism by which RRC magnitude can be increased. In general, the derived profile (Figure 3A) upregulated outward currents and downregulated inward currents. However, I_NaL_ was unexpectedly upregulated. Further analyses illuminated that I_NaL_ is important in prolonging APD_90_ to satisfy physiologic boundaries, while allowing RRC to decrease (Figures 4 and 5). This role by I_NaL_ could not be replaced by I_CaL_, another key inward current important during phases 2 and 3 of action potential repolarization. Importantly, the GB model was chosen as a platform for independent validation as it was not used in genetic algorithm training and confirmed our findings (Figure 8).

### 4.2 Relation to Prior Work on Repolarization Reserve

Most efforts made to describe repolarization reserve have quantified effects by individual ion channel currents. For example, many studies have shown that I_Ks_ can prevent extreme action potential prolongation and the development of EADs, when repolarization reserve is decreased with blocked I_Kr_ (Xiao et al., 2008; Jost et al., 2005; Varshneya et al., 2018; Silva and Rudy, 2005) or increased I_NaL_ (Printemps et al., 2019). A similar finding has also been demonstrated with I_K1_ both computationally (Ishihara et al., 2009) and in ventricular cardiomyocytes (Jost et al., 2013; Biliczki et al., 2002) with reduced repolarization reserve due to I_Kr_ block. Lastly, Nguyen et al. (Nguyen et al., 2015) found that I_to_ augmentation triggered EADs in ventricular myocytes with reduced repolarization reserve due to oxidative stress or hypokalemia. Our study expands upon such work in a few ways. First, it studies the role of nine key ion channels conductances to provide a systems-level study. Second, it utilizes an optimization approach, which removes bias in exploring ion-channels presumed to be important.

There have been studies that provide a more global view on repolarization reserve (Sarkar and Sobie, 2011; Britton et al., 2017; Miller et al., 2023). Sarkar and Sobie (2011) used multivariable regression to quantify repolarization reserve after 75% I_Kr_ block; Britton et al. (2017) used a population of models approach to study repolarization abnormalities upon multichannel ionic block of I_Kr_, I_Ks_, I_K1_, and I_CaL_; and Miller et al. (2023) published a population of models study that provided insight to ion-channel conductance correlations that play a role in LQT3 susceptibility. All these studies provide valuable insights to arrhythmia resistance in the case of decreased repolarization reserve by using ion-channel block or disease mutation models. In addition, such studies use morphology (EADs, repolarization failures, APD_90_) to measure the robustness of a cardiomyocyte rather than a quantitative metric like RRC. Therefore, this study is the first, to the best of our knowledge, to find an ion-channel conductance profile that can improve the RRC metric. It also eliminates the use of ion-channel block to reduce repolarization reserve and instead leverages bias current, which allows for all conductances to be treated and studied equally. This approach illuminated an unexpected finding by I_NaL_ (Figures 4 and 5) which suggests that using such a systems-level approach to study repolarization reserve is comprehensive and important.

### 4.3 Role of outward K^+^ currents in repolarization reserve

Both individual ion-channel (Xiao et al., 2008; Jost et al., 2005; Varshneya et al., 2018; Silva and Rudy, 2005) and systems-level (Sarkar and Sobie, 2011) repolarization reserve studies have shown that I_Ks_ increase is important to limit proarrythmic substrates when I_Kr_ is blocked. Our work agrees with these findings as the group of best GA individuals exhibit a strong negative correlation between I_Kr_ and I_Ks_ while a population of individuals with low RRC size does not (Figure S2).

This work also supports the suggestion by Ishihara et al. (2009) that I_K1_ upregulation can increase repolarization reserve (Figure 3A). I_Ks_ and I_to_ have the largest variability in parameter scaling (Figure 3A) among the best 220 individuals due to the optimization’s lack of sensitivity to these parameters (Figure 4B). The lack of sensitivity could be due to reduced density (I_Ks_), or activation early in action potential (I_to_) compared to other ion-channel conductances in the model. Specifically, I_to_ is important for repolarization in phase 1, not phase 3 and/or 4 which largely dictates repolarization reserve.

### 4.4 I_NaL_ in Repolarization Reserve

Congenital long QT syndrome type 3 (LQT3) (Moreno and Clancy, 2012), hypertrophic cardiomyopothy (HCM) (Coppini et al., 2013), hypoxia (Ju et al., 1996), and heart failure (HF) (Trenor et al., 2012; Undrovinas et al., 2006; Bai et al., 2014; Hegyi et al., 2019; Valdivia et al., 2005) are all associated with I_NaL_ enhancement and increased arrhythmogenesis. Wu et al. (Wu et al., 2006) found that proarrythmic activities and duration of the monophasic action potential increased when rabbit hearts were exposed to QT-prolonging drugs with upregulation of I_NaL_. The same group also found that I_NaL_ inhibition reversed arrhythmic activity in rabbit hearts when repolarization reserve was decreased by I_K_ inhibitors (Wu et al., 2009). Studies using single ventricular cardiomyocytes suggest I_NaL_ upregulation decreases repolarization reserve as well. For example, Sicouri et al. (Sicouri et al., 2013) concluded that I_NaL_ inhibition supressed EADs and DADs in canine cells; and Cardona et al. (Cardona et al., 2010) demonstrated *in silico* that an increase in I_NaL_ prolonged APD_90_ and led to EADs in worsened repolarization reserve conditions.

Unexpectedly, we found that an increase of I_NaL_ is important in conditions of increased repolarization reserve in two models (Figures 5 and 8). Specifically, upregulation of I_NaL_ is important for the genetic algorithm to find a physiologic ion-channel conductance profile (i.e., APD_90_ and CaT_amp_ within bounds) with increased RRC magnitude (Figure 3A). Since large increases in potassium currents abnormally shorten the action potential, an inward current is needed to appropriately balance the outward current. This highlights the importance of considering the balance of inward and outward currents as the particular role of a given ion-channel seems dependent on the repolarization reserve state, and compensatory mechanism by other ion-channel conductances in a cardiomyocyte. In neuroscience, it is established that ion-channel overlap and correlations can allow interindividual variation to be tolerated (Marder and Goaillard, 2006; Marder et al., 2007; Hudson and Prinz, 2010). This concept has been shown through *in silico* cardiomyocyte studies as well (Ballouz et al., 2021; Britton et al., 2013; Muszkiewicz et al., 2016). Most relevant, Hegyi et al. (Hegyi et al., 2020) found that APD_90_ prolongation by increased I_NaL_ is limited by I_Kr_ upregulation. They further show that this relationship is not demonstrated in models of reduced repolarization reserve (i.e., heart failure rabbit and pig cardiomyocytes). Lastly, a study by Miller et al. (Miller et al., 2023) found that susceptibility to arrhythmia is decreased with a higher I_Kr_-to-I_Na_ ratio in a LQT3 model. Therefore, in a model of increased I_Kr_, and high RRC size, increases in I_NaL_ could be important in preventing pathophysiology.

### 4.5 Cardioprotectivity of decreased I_CaL_

Like I_NaL_, I_CaL_ is a key inward current during phases 2 and 3 of the action potential and a current that APD_90_ is typically sensitive to. Unexpectedly, we demonstrated in Figure 4 that I_CaL_ was not as successful as I_NaL_ in achieving a physiologic APD_90_. To better understand this result, we attempted to rescue extreme action potential shortening in an environment of increased repolarization reserve with I_CaL_. Figure 5 demonstrates that an increase in I_CaL_ (BM-I_CaL_) decreased RRC size (−1.1 A/F) compared to BM-I_NaL_ in which APD_90_ was prolonged with I_NaL_ (−1.69 A/F). Upon further investigation we found that the large increase in I_CaL_ also increases depolarizing current by I_NCX_, which we suggest causes the reduction in RRC (Figure 6B). This was not evident when I_NaL_ was upregulated. Instead, the increase in I_NaL_ increased outward but not inward current by NCX. During diastole, NCX is in “forward” mode in which the influx of sodium and efflux of calcium generate a net inward current. During the plateau of the action potential, NCX is in “reverse” mode with opposite fluxes. Therefore, severe upregulation by NCX is not needed when I_NaL_ is increased, and consequentially, a large increase in inward current by I_NCX_ is not evident.

The downregulation of I_CaL_ appears to be important for cardioprotectivity (Figures 3, 5, and 8). We also found that I_CaL_ and I_Kr_ have a strong positive correlation (Figure 6). This agrees with a study by Ballouz et al. (Ballouz et al., 2021), which found that cells with increased coexpression of I_CaL_ and I_Kr_ at baseline generated more EADs than cells with decreased coexpression when perturbed with I_Kr_ drug block. In addition, Rees at al. completed an *in silico* and *in vitro* study to find that I_CaL_ and I_Kr_ compensate to generate a normal calcium transient in mouse ventricular cardiomyocytes. Therefore, cells with low baseline I_CaL_ that are positively correlated with I_Kr_ are more robust than uncorrelated cells with increased I_CaL_.

### 4.6 Limitations

Future directions could further improve and validate this work. First, the profile’s ability to attenuate other proarrythmic markers such as triangulation, rate dependence, and alternans could be assessed. Similar to the analysis that was completed for I_Kr_ block (Figure S3), action potential morphology between the found ion-channel conductance profile and the baseline ToR-ORd model could be compared under perturbations that trigger such proarrythmias.

Second, additional parameters could be included in GA optimization. Each individual optimized by the genetic algorithm is a set of 9 key ion-channel conductances. Including additional flux or ion-channel kinetic parameters would allow for a more global understanding of how RRC can be improved.

Third, the range over which each GA parameter varies could be increased. Each parameter was selected from a range of 33%-300% of baseline, which allowed for both healthy and potentially abnormal ionic current profiles to be included in the initial population. However, I_NaL_ and I_Na_ hit the upper and lower bounds respectively. While adding more flux or ion-channel kinetic parameters, increasing the population size, and/or running the GA optimization for more generations has the potential to add valuable information to the study, it would make the optimization problem harder, and require greater computational resources.

Fourth, this work could be validated in an additonal model system. A known limitation of electrophysiologic computational models is that there are inherent differences between them. This was highlighted in results as APD_90_ is much more sensitive to I_CaL_ in the GB model (Figure 8) than the ToR-ORd model (Figure 5). There are also known differences in ion-channel expression and dynamics between computational models and *in vitro* experiments. Therefore, it is important, but not conclusive, to validate findings in an additional model system as we did with the GB model.

Fifth, experimental feasibility could also be assessed with dynamic clamp (Ortega et al., 2018) - a method that allows for ionic currents to be artificially augmented or reduced in real time *in vitro*.

### 4.7 Conclusions

We have developed a genetic algorithm to derive a robust and cardio-protective ion-channel conductance profile, which provides a model platform to study repolarization reserve without bias or prior knowledge. Importantly, the comprehensive and systems-level approach provided unexpected mechanistic insights on how repolarization reserve could be improved. This work may have the potential to aid research in arrhythmia mitigation.

## 5 Supplemental Material

The code for the developed genetic algorithm pipeline, simulation data, and materials to reproduce manuscript figures are available at: https://github.com/Christini-Lab/rrc-conductance-profile.git

## 7 Author Contributions

K.E.F. designed research, performed simulations and computations, analyzed data, prepared figures, and drafted manuscript. K.E.F., A.P.C., T.K.M., and D.J.C. interpreted results. T.K.M., A.P.C. and D.J.C. edited and revised manuscript. K.E.F., A.P.C., T.K.M., and D.J.C. approved final version of manuscript.

## References

Bai J, Wang K, Bai X, Yuan Y & Zhang H (2014). Pro-arrhythmic effects of increased late sodium current in failing human heart. Computing in Cardiology 2014 pp. 857–860.

Ballouz S, Mangala MM, Perry MD, Heitmann S, Gillis JA, Hill AP & Vandenberg JI (2021). Co-expression of calcium and hERG potassium channels reduces the incidence of proarrhythmic events. Cardiovascular Research 117, 2216–2227.

Behr E & Ensam B (2016). New approaches to predicting the risk of sudden death. Clinical Medicine 16, 283–283.

Beuckelmann DJ, Näbauer M & Erdmann E (1992). Intracellular calcium handling in isolated ventricular myocytes from patients with terminal heart failure. Circulation 85, 1046–1055.

Biliczki P, Virág L, Iost N, Papp JG & Varró A (2002). Interaction of different potassium channels in cardiac repolarization in dog ventricular preparations: role of repolarization reserve. British Journal of Pharmacology 137, 361–368.

Bot C, Kherlopian A, Ortega F, Christini D & Krogh-Madsen T (2012). Rapid Genetic Algorithm Optimization of a Mouse Computational Model: Benefits for Anthropomorphization of Neonatal Mouse Cardiomyocytes. Frontiers in Physiology 3, 421.

Britton OJ, Bueno-Orovio A, Van Ammel K, Lu HR, Towart R, Gallacher DJ & Rodriguez B (2013). Experimentally calibrated population of models predicts and explains intersubject variability in cardiac cellular electrophysiology. Proceedings of the National Academy of Sciences 110, E2098–E2105.

Britton OJ, Bueno-Orovio A, Virág L, Varró A & Rodriguez B (2017). The Electrogenic Na+/K+ Pump Is a Key Determinant of Repolarization Abnormality Susceptibility in Human Ventricular Cardiomyocytes: A Population-Based Simulation Study. Front Physiol 8, 278.

Cardona K, Trenor B, Romero L, Ferrero J & Saiz J (2010). Role of the late sodium current in arrhythmias related to low repolarization reserve pp. 617–620.

Clark AP, Wei S, Kalola D, Krogh-Madsen T & Christini DJ (2022). An in silico–in vitro pipeline for drug cardiotoxicity screening identifies ionic pro-arrhythmia mechanisms. British Journal of Pharmacology 179, 4829–4843.

Clerx M, Collins P, de Lange E & Volders PGA (2016). Myokit: A simple interface to cardiac cellular electrophysiology. Progress in biophysics and molecular biology 120, 100–14.

Coppini R, Ferrantini C, Yao L, Fan P, Del Lungo M, Stillitano F, Sartiani L, Tosi B, Suffredini S, Tesi C, Yacoub M, Olivotto I, Belardinelli L, Poggesi C, Cerbai E & Mugelli A (2013). Late Sodium Current Inhibition Reverses Electromechanical Dysfunction in Human Hypertrophic Cardiomyopathy. Circulation 127, 575–584.

Fenichel RR, Malik M, Antzelevitch C, Sanguinetti M, Roden DM, Priori SG, Ruskin JN, Lipicky RJ, Cantilena LR & Force IAT (2004). Drug-Induced Torsades de Pointes and Implications for Drug Development. Journal of Cardiovascular Electrophysiology 15, 475–495.

Fortin FA, De Rainville FM, Gardner MAG, Parizeau M & Gagné C (2012). DEAP: evolutionary algorithms made easy. J. Mach. Learn. Res. 13, 2171–2175.

Gaur N, Ortega F, Verkerk AO, Mengarelli I, Krogh-Madsen T, Christini DJ, Coronel R & Vigmond EJ (2020). Validation of quantitative measure of repolarization reserve as a novel marker of drug induced proarrhythmia. Journal of Molecular and Cellular Cardiology 145, 122–132.

Grandi E, Pasqualini FS & Bers DM (2010). A novel computational model of the human ventricular action potential and Ca transient. Journal of Molecular and Cellular Cardiology 48, 112–121.

Groenendaal W, Ortega FA, Kherlopian AR, Zygmunt AC, Krogh-Madsen T & Christini DJ (2015). Cell-Specific Cardiac Electrophysiology Models. PLOS Computational Biology 11, e1004242.

Hancox JC, McPate MJ, E. Harchi A & Zhang Yh (2008). The hERG potassium channel and hERG screening for drug-induced torsades de pointes. Pharmacology & Therapeutics 119, 118–132.

Hayashi M, Shimizu W & Albert CM (2015). The Spectrum of Epidemiology Underlying Sudden Cardiac Death. Circ Res 116, 1887–1906.

Hegyi B, Chen-Izu Y, Izu LT, Rajamani S, Belardinelli L, Bers DM & Bányász T (2020). Balance Between Rapid Delayed Rectifier K+ Current and Late Na+ Current on Ventricular Repolarization. Circulation: Arrhythmia and Electrophysiology 13, e008130.

Hegyi B, Morotti S, Liu C, Ginsburg KS, Bossuyt J, Belardinelli L, Izu LT, Chen-Izu Y, Bányász T, Grandi E & Bers DM (2019). Enhanced Depolarization Drive in Failing Rabbit Ventricular Myocytes. Circulation: Arrhythmia and Electrophysiology 12, e007061.

Hudson AE & Prinz AA (2010). Conductance Ratios and Cellular Identity. PLOS Computational Biology 6, e1000838.

Ishihara K, Sarai N, Asakura K, Noma A & Matsuoka S (2009). Role of Mg2+ block of the inward rectifier K+ current in cardiac repolarization reserve: A quantitative simulation. Journal of Molecular and Cellular Cardiology 47, 76–84.

Jost N, Virág L, Bitay M, Takács J, Lengyel C, Biliczki P, Nagy Z, Bogáts G, Lathrop DA, Papp JG & Varró A (2005). Restricting Excessive Cardiac Action Potential and QT Prolongation. Circulation 112, 1392–1399.

Jost N, Virág L, Comtois P, Ördög B, Szuts V, Seprényi G, Bitay M, Kohajda Z, Koncz I, Nagy N, Szél T, Magyar J, Kovács M, Puskás LG, Lengyel C, Wettwer E, Ravens U, Nánási PP, Papp JG, Varró A & Nattel S (2013). Ionic mechanisms limiting cardiac repolarization reserve in humans compared to dogs. The Journal of Physiology 591, 4189–4206.

Ju YK, Saint DA & Gage PW (1996). Hypoxia increases persistent sodium current in rat ventricular myocytes. J Physiol 497, 337–347.

Kannankeril P, Roden DM & Darbar D (2010). Drug-Induced Long QT Syndrome. Pharmacol Rev 62, 760–781.

Kherlopian AR, Ortega FA & Christini DJ (2011). Cardiac myocyte model parameter sensitivity analysis and model transformation using a genetic algorithm. Association for Computing Machinery 11, 755–758.

Krogh-Madsen T, Jacobson AF, Ortega FA & Christini DJ (2017). Global Optimization of Ventricular Myocyte Model to Multi-Variable Objective Improves Predictions of Drug-Induced Torsades de Pointes. Front Physiol 8, 1059.

Marder E & Goaillard JM (2006). Variability, compensation and homeostasis in neuron and network function. Nat Rev Neurosci 7, 563–574.

Marder E, Tobin AE & Grashow R (2007). How tightly tuned are network parameters? Insight from computational and experimental studies in small rhythmic motor networks. Prog Brain Res 165, 193–200.

Miller JA, Moise N & Weinberg SH (2023). Modeling incomplete penetrance in long QT syndrome type 3 through ion channel heterogeneity: an in silico population study. Am J Physiol Heart Circ Physiol 324, H179–H197.

Moreno JD & Clancy CE (2012). Pathophysiology of the cardiac late Na current and its potential as a drug target. Journal of Molecular and Cellular Cardiology 52, 608–619.

Muszkiewicz A, Britton OJ, Gemmell P, Passini E, Sánchez C, Zhou X, Carusi A, Quinn TA, Burrage K, Bueno-Orovio A & Rodriguez B (2016). Variability in cardiac electrophysiology: Using experimentally-calibrated populations of models to move beyond the single virtual physiological human paradigm. Progress in Biophysics and Molecular Biology 120, 115–127.

Nguyen TP, Singh N, Xie Y, Qu Z & Weiss JN (2015). Repolarization Reserve Evolves Dynamically During the Cardiac Action Potential. Circulation: Arrhythmia and Electrophysiology 8, 694–702.

O’Hara T, Virág L, Varró A & Rudy Y (2011). Simulation of the Undiseased Human Cardiac Ventricular Action Potential: Model Formulation and Experimental Validation. PLOS Computational Biology 7, e1002061.

Ortega FA, Grandi E, Krogh-Madsen T & Christini DJ (2018). Applications of Dynamic Clamp to Cardiac Arrhythmia Research: Role in Drug Target Discovery and Safety Pharmacology Testing. Frontiers in Physiology 8, 1099.

Passini E, Britton OJ, Lu HR, Rohrbacher J, Hermans AN, Gallacher DJ, Greig RJH, Bueno-Orovio A & Rodriguez B (2017). Human In Silico Drug Trials Demonstrate Higher Accuracy than Animal Models in Predicting Clinical Pro-Arrhythmic Cardiotoxicity. Frontiers in Physiology 8, 668.

Printemps R, Salvetat C, Faivre JF, Grand M, Bois P & Moha ou Maati H (2019). Role of Cardiac IKs Current in Repolarization Reserve Process During Late Sodium Current (INaL) Activation. Cardiology and Cardiovascular Medicine 03, 168–185.

Roden DM (2006). Long QT syndrome: reduced repolarization reserve and the genetic link. Journal of Internal Medicine 259, 59–69.

Roden DM (1998). Taking the “Idio” out of “Idiosyncratic”: Predicting Torsades de Pointes. Pacing and Clinical Electrophysiology 21, 1029–1034.

Sarkar AX & Sobie EA (2011). Quantification of repolarization reserve to understand interpatient variability in the response to proarrhythmic drugs: A computational analysis. Heart Rhythm 8, 1749–1755.

Sicouri S, Belardinelli L & Antzelevitch C (2013). Antiarrhythmic effects of the highly selective late sodium channel current blocker GS-458967. Heart Rhythm 10, 1036–1043.

Silva J & Rudy Y (2005). Subunit Interaction Determines IKs Participation in Cardiac Repolarization and Repolarization Reserve. Circulation 112, 1384–1391.

Syed Z, Vigmond E, Nattel S & Leon LJ (2005). Atrial cell action potential parameter fitting using genetic algorithms. Med. Biol. Eng. Comput. 43, 561–571.

Tomek J, Bueno-Orovio A, Passini E, Zhou X, Minchole A, Britton O, Bartolucci C, Severi S, Shrier A, Virag L, Varro A & Rodriguez B (2019). Development, calibration, and validation of a novel human ventricular myocyte model in health, disease, and drug block. eLife 8, e48890.

Trenor B, Cardona K, Gomez JF, Rajamani S, Jr JMF, Belardinelli L & Saiz J (2012). Simulation and Mechanistic Investigation of the Arrhythmogenic Role of the Late Sodium Current in Human Heart Failure. PloS One 7(3), e32659.

Undrovinas AI, Belardinelli L, Undrovinas NA & Sabbah HN (2006). Ranolazine Improves Abnormal Repolarization and Contraction in Left Ventricular Myocytes of Dogs with Heart Failure by Inhibiting Late Sodium Current. Journal of Cardiovascular Electrophysiology 17, S169–S177.

Valdivia CR, Chu WW, Pu J, Foell JD, Haworth RA, Wolff MR, Kamp TJ & Makielski JC (2005). Increased late sodium current in myocytes from a canine heart failure model and from failing human heart. Journal of Molecular and Cellular Cardiology 38, 475–483.

Varró A & Baczkó I (2011). Cardiac ventricular repolarization reserve: a principle for understanding drug-related proarrhythmic risk. British Journal of Pharmacology 164, 14–36.

Varshneya M, Devenyi RA & Sobie EA (2018). Slow Delayed Rectifier Current Protects Ventricular Myocytes From Arrhythmic Dynamics Across Multiple Species. Circulation: Arrhythmia and Electrophysiology 11, e006558.

Weiss JN, Karma A, MacLellan WR, Deng M, Rau CD, Rees CM, Wang J, Wisniewski N, Eskin E, Horvath S, Qu Z, Wang Y & Lusis AJ (2012). “Good Enough Solutions” and the Genetics of Complex Diseases. Circulation Research 111, 493–504.

Wu L, Rajamani S, Li H, January CT, Shryock JC & Belardinelli L (2009). Reduction of repolarization reserve unmasks the proarrhythmic role of endogenous late Na+ current in the heart. American Journal of Physiology-Heart and Circulatory Physiology 297, H1048–H1057.

Wu L, Shryock JC, Song Y & Belardinelli L (2006). An Increase in Late Sodium Current Potentiates the Proarrhythmic Activities of Low-Risk QT-Prolonging Drugs in Female Rabbit Hearts. J Pharmacol Exp Ther 316, 718–726.

Xiao L, Xiao J, Luo X, Lin H, Wang Z & Nattel S (2008). Feedback Remodeling of Cardiac Potassium Current Expression. Circulation 118, 983–992.

