## Supplemental Material for "Optimization of a Cardiomyocyte Model Illuminates Role of Increased I_NaL_ in Repolarization Reserve"

### 1 Model Updates

Since the GB model does not include late sodium, the $I_{NaL}$ equations were extracted from the ToR-ORd model source code and added to the GB model. The late sodium conductance parameter was scaled by the ratio of peak sodium between both models (356 A/F (GB) / 294 A/F (ToR-ORd)). Lastly, CaMKII phosphorylation was added to the GB model as well since the ToR-ORd $I_{NaL}$ formulation is dependent on it.

#### 1.1 Definitions and Abbreviations

| parameter | model code variable | definition |
|---|---|---|
| $m_{L,\infty}$ | mLss | steady state activation for $I_{NaL}$ |
| $\tau_{m,L}$ | tmL | time constant of gate m |
| $m_L$ | mL | activation for $I_{NaL}$ |
| $h_{L,\infty}$ | hLss | steady state inactivation for $I_{NaL}$ |
| $\tau_{h,L}$ | thL | time constant of gate h |
| $h_L$ | hL | inactivation for $I_{NaL}$ |
| $h_{L,CaMK,\infty}$ | hLssp | steady state inactivation for CaMK phosphorylated $I_{NaL}$ |
| $\tau_{h,LCaMK}$ | thLp | time constant of h gate phosphorylated |
| $h_{L,CaMK}$ | hLp | inactivation for CaMK phosphorylated $I_{NaL}$ |
| $\phi_{INaL,CaMK}$ | fINaLp | fraction of $I_{NaL}$ channels phosphorylated by CaMK |
| $G_{NaL}$ | GNaL | maximum conductance of INaL |
| $I_{NaL,junc}$ | INaL_junc | current through $I_{NaL}$ channels in the junctional space |
| $I_{NaL,sl}$ | INaL_sl | current through $I_{NaL}$ channels in the subsarcolema space |
| $I_{NaL}$ | INaL | current through $I_{NaL}$ channels |
| $\alpha_{CaMK}$ | aCaMK | phosphorylation rates of CaMK |
| $\beta_{CaMK}$ | bCaMK | dephosphorylation rates of CaMK |
| $CaMK_0$ | CaMKo | fraction of active CaMK binding sites at equilibrium |
| $CaMK_{active}$ | CaMKa | fraction of active CaMK binding sites |
| $CaMK_{bound}$ | CaMKb | fraction of CaMK binding sites bound to $Ca^{2+}$/calmodulin |
| $CaMK_{trap}$ | CaMKt | fraction of autonomous CaMK binding sites with trapped calmodulin |

#### 1.2 Initial condition additions

$$m_L = 0.0001629$$

$$h_L = 0.5255$$

$$h_{L,CaMK} = 0.2872$$

$$CaMK_{trap} = 0.0111$$

##### 1.3 $\mathbf{I_{NaL}}$

$$m_{L,\infty} = \frac{1}{1 + e^{\frac{-V-42.85}{5.264}}}$$

$$\tau_{m,L} = 0.1292 * e^{-(\frac{V+45.79}{15.54})^2} + 0.06487 * e^{-(\frac{V-4.823}{51.12})^2}$$

$$\frac{d(m_L)}{dt} = \frac{m_{L,\infty} - m_L}{\tau_{m,L}}$$

$$h_{L,\infty} = \frac{1}{1 + e^{\frac{V+87.61}{7.488}}}$$

$$\tau_{h,L} = 200ms$$

$$\frac{d(h_L)}{dt} = \frac{h_{L,\infty} - h_L}{t_{h,L}}$$

$$h_{L,CaMK,\infty} = \frac{1}{1 + e^{\frac{V+93.81}{7.488}}}$$

$$\tau_{h,LCaMK} = 3 * \tau_{h,L}$$

$$\frac{d(h_{L,CaMK})}{dt} = \frac{h_{L,CaMK,\infty} - h_{L,CaMK}}{\tau_{h,LCaMK}}$$

$$\phi_{INaL,CaMK} = \frac{1}{1 + \frac{0.15}{CaMK_{active}}}$$

$$G_{NaL} = 0.029 * \frac{356}{294} \frac{mS}{\mu F}$$

$$I_{NaL,junc} = G_{NaL} * (V - E_{Na_{junc}}) * m_L * ((1 - \phi_{INaL,CaMK}) * h_L + \phi_{INaL,CaMK} * h_{L,CaMK})$$

$$I_{NaL,sl} = G_{NaL} * (V - E_{Na_{sl}}) * m_L * ((1 - \phi_{INaL,CaMK}) * h_L + \phi_{INaL,CaMK} * h_{L,CaMK})$$

$$I_{NaL} = I_{NaL,junc} + I_{NaL,sl}$$

where $E_{Na_{junc}}$ and $E_{Na_{sl}}$ are the Nerst potentials for sodium in the junctional and subsarcolemmal compartments, respectively. The equations for these two parameters were not changed in respect to the baseline BM model and can be found in the supplmetary material of Grandi et al. [**grandi˙novel˙2010**].

##### 1.4 CaMK

$$\alpha_{CaMK} = 0.05 \text{ ms}^{-1}$$

$$\beta_{CaMK} = 0.00068 \text{ ms}^{-1}$$

$$CaMK_0 = 0.05$$

$$CaMK_{bound} = CaMK_0 * \frac{1 - CaMK_{trap}}{1 + \frac{0.0015}{Ca_i}}$$

$$CaMK_{active} = CaMK_{bound} + CaMK_{trap}$$

$$\frac{d(CaMK_{trap})}{dt} = \alpha_{CaMK} * CaMK_{bound} * (CaMK_{bound} + CaMK_{trap}) - \beta_{CaMK} * CaMK_{trap}$$

where $Ca_i$ is intracellular calcium which was not changed in respect to the baseline GB model.

#### 2 Supplemental Figures

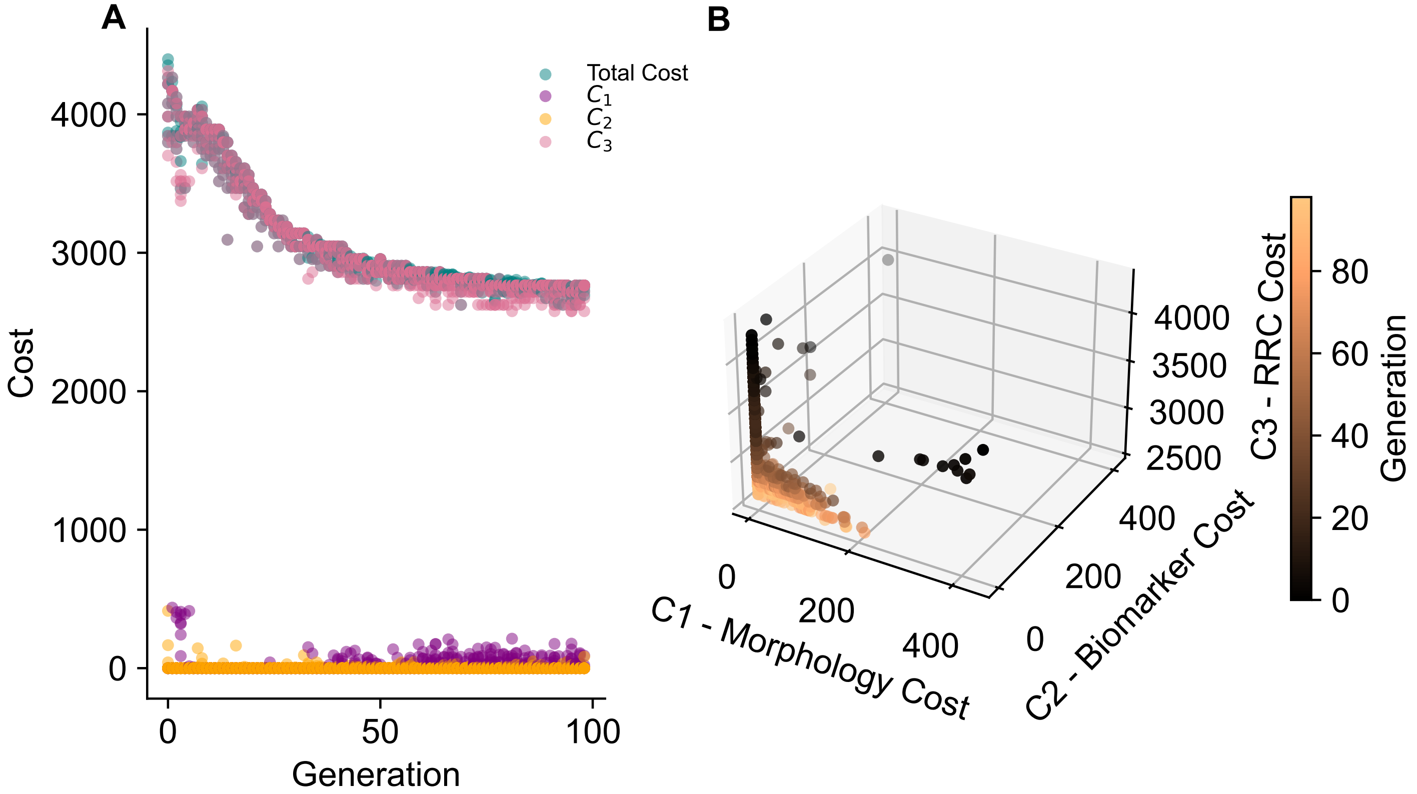


**Figure S 1: Evolution of the best individuals in GA optimization and their respective cost components.** A. The total cost of the best individual at each generation and its respective components ($C_1$, $C_2$, and $C_3$) are plotted for all 8 GA runs. Therefore, 32 (4 cost terms x the best individual from all 8 GA trials) points are plotted for each generation. B. The best individual in each generation from all 8 GA trials plotted by its $C_1$, $C_2$, and $C_3$ values which sum to calculate the total cost. Each point is colored by generation.

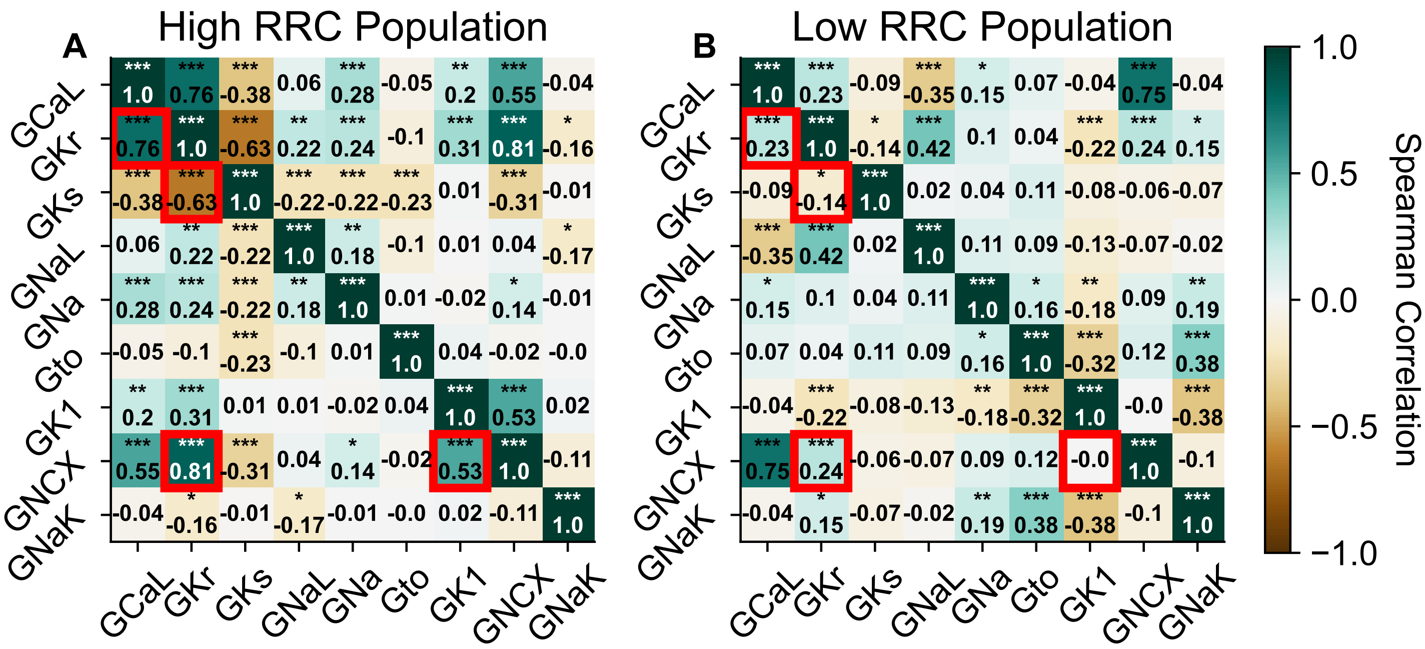


**Figure S 2: Correlations between specific ion-channels are present in a group of cells with high RRC size but not in a group with low RRC.** Spearman correlations between all nine ion-channel profile conductances for a population of models with A. high and B. low RRC magnitude. The high RRC population is the 220 best individuals present in all generations of all 8 GA trials. These individuals all have a $C_3$ error below 2800. The population of models with low RRC represent the 233 worst individuals present in all generations of all 8 GA trials. These individuals all have a $C_3$ error greater than 5200. The red box highlights strong correlations in A that are not present in B.

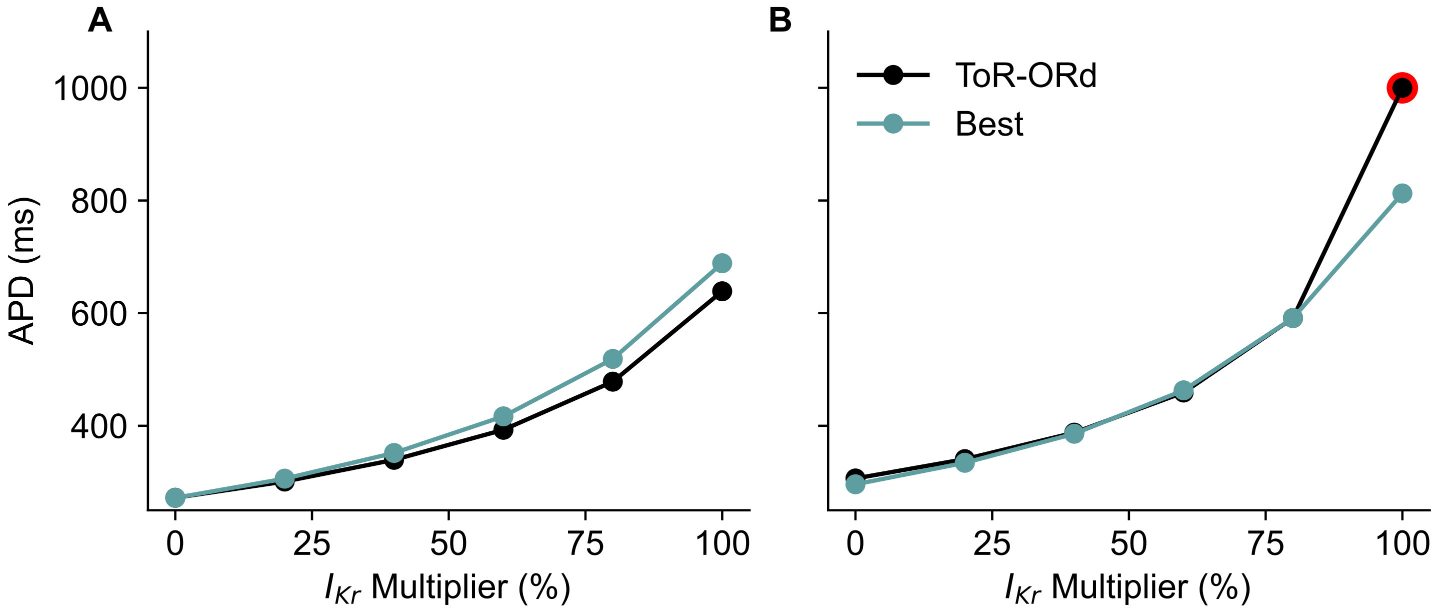


**Figure S 3: Evaluation of APD after $I_K$ perturbations.** A. APD value after 0, 20%, 40%, 60%, 80%, and 99% IKr block. B. The same analysis as A was repeated but with an additional block of the background potassium channel ($I_{Kb}$). The perturbations were applied to both the baseline ToR-ORd model (black) and a representative optimized model. The red circle represents a simulation with a repolarization abnormality.
